# Enhancing CAR-T Cell Metabolism to Overcome Hypoxic Conditions in the Brain Tumor Microenvironment

**DOI:** 10.1101/2023.11.13.566775

**Authors:** Ryusuke Hatae, Keith Kyewalabye, Akane Yamamichi, Tiffany Chen, Su Phyu, Pavlina Chuntova, Takahide Nejo, Lauren S. Levine, Matthew H. Spitzer, Hideho Okada

## Abstract

The efficacy of chimeric antigen receptor (CAR)-T therapy has been limited against brain tumors to date. CAR-T cells infiltrating syngeneic intracerebral SB28-EGFRvIII glioma revealed impaired mitochondrial ATP production and a markedly hypoxic status compared to ones migrating to subcutaneous tumors. Drug screenings to improve metabolic states of T cells under hypoxic conditions led us to evaluate the combination of AMPK activator Metformin and the mTOR inhibitor Rapamycin (Met+Rap). Met+Rap-pretreated mouse CAR-T cells showed activated PPAR-gamma coactivator 1α (PGC-1α) through mTOR inhibition and AMPK activation, and a higher level of mitochondrial spare respiratory capacity than those pretreated with individual drugs or without pretreatment. Moreover, Met+Rap-pretreated CAR-T cells demonstrated persistent and effective anti-glioma cytotoxic activities in the hypoxic condition. Furthermore, a single intravenous infusion of Met+Rap-pretreated CAR-T cells significantly extended the survival of mice bearing intracerebral SB28-EGFRvIII gliomas. Mass cytometric analyses highlighted increased glioma-infiltrating CAR-T cells in the Met+Rap group with fewer Ly6c+ CD11b+ monocytic myeloid-derived suppressor cells in the tumors. Finally, human CAR-T cells pretreated with Met+Rap recapitulated the observations with murine CAR-T cells, demonstrating improved functions in vitro hypoxic conditions. These findings advocate for translational and clinical exploration of Met+Rap-pretreated CAR-T cells in human trials.

## Introduction

Glioblastoma multiforme (GBM) is an aggressive and lethal brain tumor despite the combined standard of care with maximum surgical resection, radiation, and chemotherapy. One potential approach to treating GBM is immunotherapy; however, despite promising results in several other types of cancer, immunotherapy has not been effective against GBM (1). One of the key challenges to successful immunotherapy in GBM is the highly immunosuppressive tumor microenvironment characterized by a number of mechanisms (2), including hypoxic conditions (3). As such, recent research has focused on developing innovative strategies to overcome these challenges and improve the effectiveness of immunotherapy (4, 5).

Chimeric antigen receptor T-cell (CAR-T) therapy has demonstrated significant efficacy against hematologic malignancies (6). However, its therapeutic potential remains limited against solid tumors, including brain neoplasms (5). The metabolic status of immune cells has recently been recognized as a critical factor in cancer immunotherapy. Glycolytic metabolism is essential for effector T cells, and oxidative phosphorylation (OXPHOS), which occurs in mitochondria, is crucial for the high survival capability of memory T cells (7). Furthermore, glycolysis and OXPHOS are known to be decreased in exhausted T cells (8). In the tumor microenvironment, hypoxic conditions and chronic antigen stimulation rapidly diminish T cell mitochondrial function and lead to exhaustion (9). Therefore, we hypothesized that enhancing the mitochondrial function of CAR-T cells could prevent them from an exhaustion status in the hypoxic microenvironment of GBM. To address the hypothesis, in this study, we investigated the preconditioning of CAR-T cells with metabolic regulators before infusion and examined its translational potential.

## Results

### CAR-T cells lose OXPHOS activity in the glioma microenvironment

To investigate the metabolic status within the tumor microenvironment, we leveraged the experimental system of anti-EGFRvIII-CAR-T cells isolated from EGFRvIII-CAR-transgenic mice that we established previously (10). This system allows us to obtain a large number of CAR-T cells with consistent quality, thus enabling us to perform extensive experiments with high scientific rigor. We evaluated the metabolic status of CAR-T cells following intravenous (IV) infusion into syngeneic C57BL/6J mice bearing intracerebral SB28 tumors expressing human EGFRVIII (SB28 hEGFRvIII). The mice received a single IV infusion of 3×10^6^ anti-EGFRvIII CAR-T cells. A cohort of 3 mice was sacrificed every 3 days post-infusion up to 21 days, the glioma tissues were harvested, and the metabolic status of brain tumor-infiltrating CAR-T cells was evaluated by flow cytometry (Figure 1A, top). The expression levels of a glycolytic marker glucose transporter 1 (Glut1) were not significantly different between the CAR-T cells isolated from the glioma tissues and those derived from the spleen (Figure 1A, bottom). On the other hand, the expression levels of ATP synthase (ATP5a), a marker of OXPHOS, in the glioma-infiltrating CAR-T cells continuously decreased over time, while those in the spleen-derived CAR-T cells did not show such decreasing trends (Figure 1A, bottom and Fig 1B).

**Figure 1.**
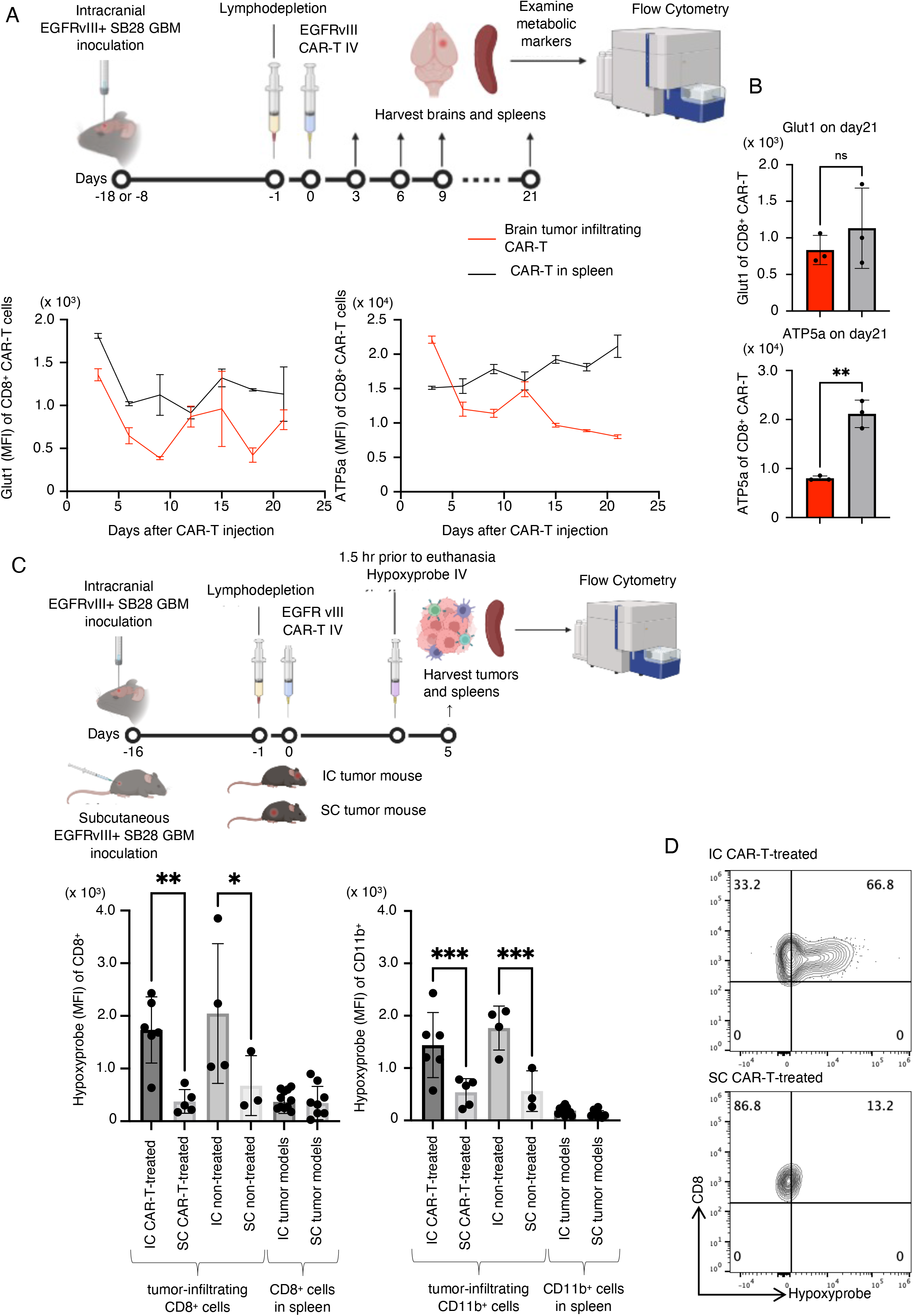
Exhaustion of CAR-T cells associated with reduced OXPHOS activity in the hypoxic glioma microenvironment. (A) The experimental design to evaluate glioma-infiltrating CAR-T cells (upper panel). Longitudinal changes in Glut1 and ATP5a, markers of the glycolytic system and OXPHOS, respectively, in glioma-infiltrating CD8+ CAR-T cells (Lower panels). (B) Expression of Glut1(top) and ATP5a (bottom) by mean fluorescence intensity (MFI) in CD8+ CAR-T cells extracted from the spleen (grey) and tumor (red) on Day 21. (C) The design for analyzing hypoxic conditions in vivo (upper panel). Uptake of hypoxyprobe by CD8+ (left) or CD11b+ (right) leukocytes infiltrating intracranial tumor models (IC) or subcutaneous tumor models (SC) SB28 mEGFRvIII gliomas or spleens of glioma-bearing mice. (D) Representative histograms (IC or SC tumors in CAR-treated mice) showing the positive staining with hypoxyprobe on CD8+ BIL but not on CD8+ CAR-T cells isolated from SC tumors. Error bars show the mean with SD. **P < 0.01 by Mann-Whitney test (B). *P < 0.05; **P < 0.01; ***P <0.001 by one-way ANOVA analysis followed by Tukey’s multiple comparison test (C).

### The brain tumor microenvironment is hypoxic

Since OXPHOS is a cellular process that produces a large amount of ATP by utilizing oxygen in the intracellular mitochondria, we hypothesized that the decreased OXPHOS activity in CAR-T cells might be due to the hypoxic glioma microenvironment. To address this hypothesis, in mice bearing day 16 SB28 tumors expressing murine EGFRvIII (SB28 mEGFRvIII) in the brain or subcutaneous space in the right flank of syngeneic C57BL/6J mice, we IV administered anti-EGFRvIII CAR-T cells. We used SB28 mEGFRvIII in our in vivo studies because of more consistent Luciferase expression in the mEGFRvIII version than SB28 hEGFRvIII (data not shown). Five days later, hypoxyprobe, an agent specifically taken up by hypoxic cells, was administered to the mice, followed by harvesting of the glioma tissues and spleens at 1.5 hours after the hypoxyprobe infusion (Figure 1C, top). Flow cytometric evaluations of CD8^+^ and CD11b^+^ cells isolated from these organs revealed that both CD8^+^ and CD11b^+^ populations in the intracerebral glioma tissue were significantly more hypoxic than those in subcutaneous tumors regardless of CAR-T cell administration Figure 1C, bottom and 1D).

### Pretreatment of CAR-T cells with Metformin and Rapamycin improves the sustained function of CAR-T cells in hypoxic conditions

We then hypothesized that treating CAR-T cells with mitochondria-activating drugs before infusion ex vivo would improve the function of CAR-T cells in hypoxic conditions. In vitro, under the normoxic condition (Figure 2 A, left), CAR-T cells maintained their potent cytotoxic function against the glioma cells, even after new glioma cells were introduced every two days for three cycles. On the other hand, under the hypoxic condition, CAR-T cells lost their antitumor effect after three cycles of tumor cell challenge (Figure 2 A, right). To develop a strategy to overcome reduced cytotoxicity in hypoxic conditions, we selected and tested Metformin (Met), Rapamycin (Rap), 2-Deoxyglucose (2DG), and Dichloroacetate (DCA) because these have been studied in combination with PD-1 inhibitory antibodies and are expected to activate mitochondria (Table 1) (11). We treated CAR-T cells with each of these OXPHOS-activating metabolic regulators prior to co-culturing them with SB28 mEGFRvIII tumor cells to evaluate whether the antitumor effect could improve under the hypoxic condition (Figure 2B, top). Among the drugs evaluated, only the combination of Met and Rap (Met+Rap) enhanced the persistent antitumor effect of CAR-T cells under the hypoxic condition (Figure 2B, bottom).

**Figure 2.**
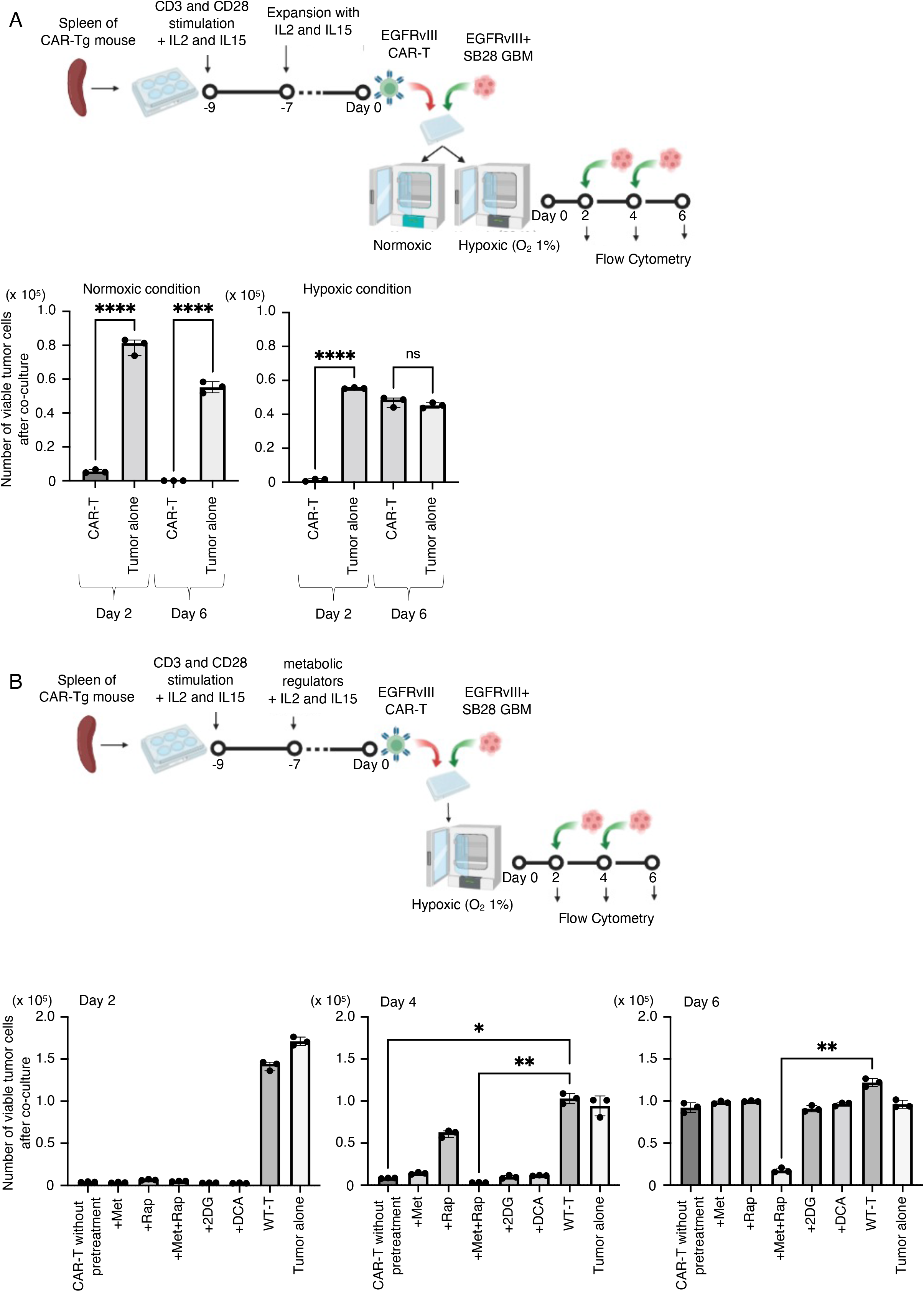
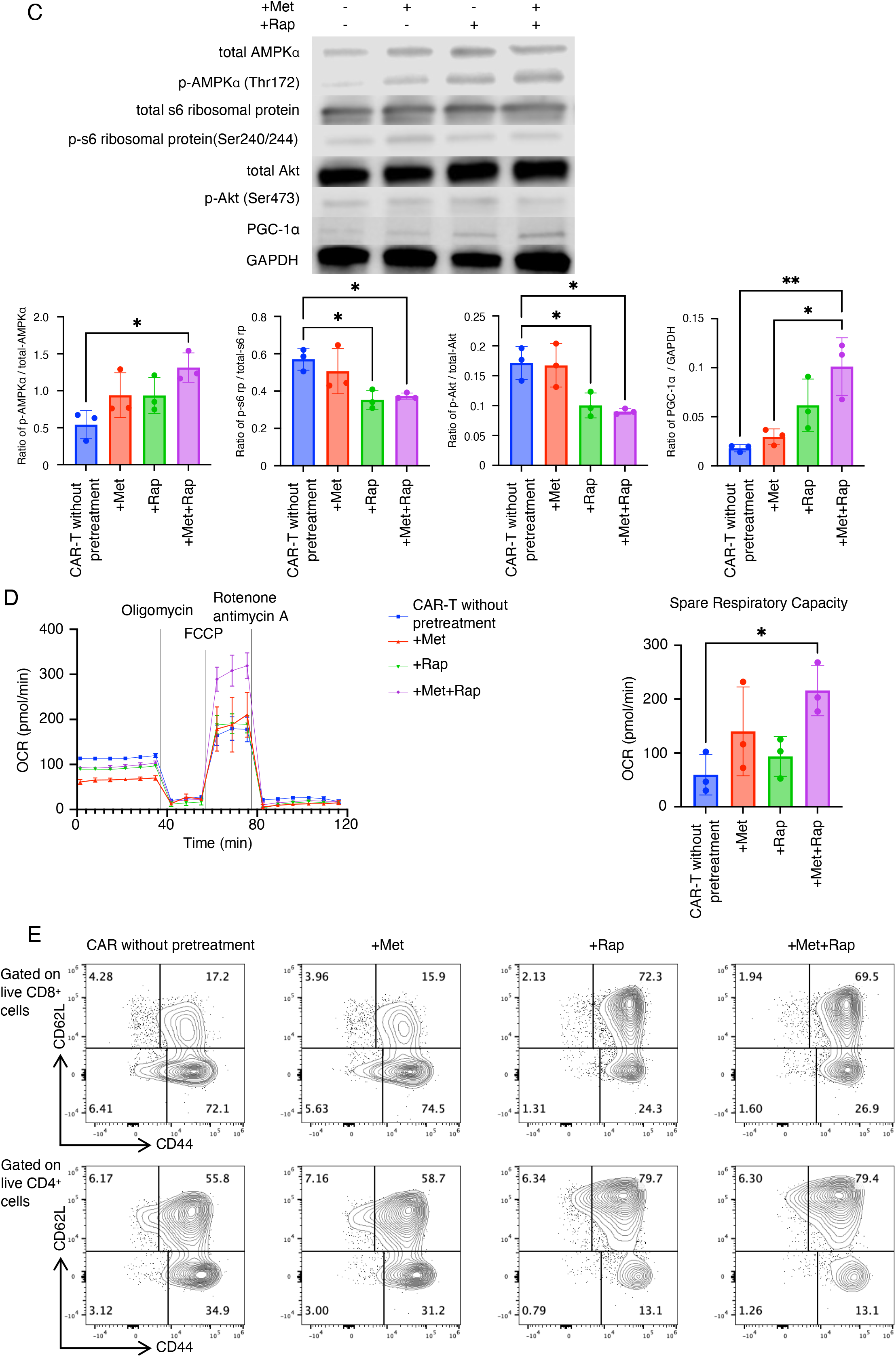
A combination of Metformin and Rapamycin (Met+Rap) promotes the persistent function of CAR-T cells in hypoxic condition. (A) The design of the co-culture experiment (upper panel). The number of viable tumor cells after co-culture under normoxic condition (left) or hypoxic condition (right). (B) The design for evaluating metabolic regulators (upper panel). The number of viable tumor cells on Day 2 (left), Day 4 (middle), and Day 6 (right). (C) Representative immunoblot of phosphorylation of AMPK ɑ, s6 ribosomal protein and Akt and the expression of total AMPK ɑ, total s6 ribosomal protein, total Akt and PGC-1ɑ in pretreated CD8+ CAR-T cells. The bar charts represent quantitative comparisons between the groups (n = 3/group). (D) OCR of CAR-T cells was measured by Seahorse XFe96 analyzer on Day 0 (left). Data represent the means ± SEM. Spare Respiratory Capacity levels were calculated (right). (E) Flow cytometric plots for CD44 and CD62L in CD8+ and CD4+ CAR-T cells on Day 0. Error bars show the mean with SD. *p < 0.05, **p < 0.01, ****p < 0.0001 by one-way ANOVA test followed by Tukey’s multiple comparisons test.

**Table 1.** List of metabolic regulators used in the drug screening.

| Compound name | Description | Vendor | Dose |
| --- | --- | --- | --- |
| Metformin (Met) | AMPK activator | Sigma-Aldrich | 10uM |
| Rapamycin (Rap) | mTORC1 inhibitor | Sigma-Aldrich | 20nM |
| 2-Deoxy-D-glucose<br>(2DG) | Glycolysis inhibitor | Sigma-Aldrich | 0.5mM |
| Dichloroacetic acid<br>(DCA) | Mitochondrial pyruvate<br>dehydrogenase kinase inhibitor | Fisher Scientific | 5mM |

### Ex vivo Met+Rap pretreatment enhances the mitochondrial respiratory capacity by activating AMPK and PGC-1ɑ in CAR-T cells

To unravel the mechanism underlying the effect of Met+Rap on CAR-T cells, we investigated the activation of AMPK and mTOR pathways in CD8^+^ CAR-T cells after treatment with Met or Rap alone, or the combination of Met+Rap. The Met+Rap treatment activated AMPK and inhibited phospho-S6 ribosomal protein and phospho-Akt ser473 (downstream targets of mammalian target of rapamycin complex (mTORC) 1 and mTORC2, respectively) (Figure 2C). Furthermore, Met+Rap pretreatment led to the upregulation of PPAR-gamma coactivator 1α (PGC-1α), the master regulator of mitochondrial biogenesis and function (12), likely due to the combined effects of Met-induced AMPK activation and Rap-induced mTOR inhibition. It has been reported that T cells overexpressing PGC1ɑ are less prone to exhaustion in the tumor microenvironment (13).

Therefore, based on the upregulation of PGC1ɑ, we hypothesized that Met+Rap combination treatment would improve the mitochondrial energy metabolism of CAR-T-cells, leading to better persistency and antitumor cytotoxicity even under hypoxic conditions. To address this, we conducted the Seahorse assay-based Mito stress test with CAR-T cells pretreated with metabolic regulators and found that Met+Rap treatment significantly increased the spare respiratory capacity (SRC) of CAR-T cells (Figure 2D). This observation was encouraging because previous studies have shown that cells with a higher SRC can survive better under hypoxic conditions (14). Importantly, these findings were consistently observed in CAR-T cells isolated from both male and female mice (Supplementary Figure 1 A-C).

Several publications reported the ability of Met or Rap to enhance the central memory T cell population (15, 16). Consistent with these findings, our data revealed that CAR-T cells pretreated with Rap or Met+Rap demonstrated a high percentage of central memory T cells (Figure 2E). In contrast, those pretreated with Met alone failed to show an increase of this population. Memory T cells generally exhibit a higher mitochondrial SRC (17). While the Rap-and Met+Rap pretreatments gave rise to similar numbers of central memory population, the Met+Rap pretreatment augmented the SRC relative to Rap alone, suggesting a more pronounced mitochondrial activation induced by the combined Met+Rap pretreatment (Figure 2D and Supplementary Figure 1C).

### Metabolic regulator therapy promotes the resistance of CAR-T cells against exhaustion

To investigate whether Met+Rap treatment would save CAR-T cells from exhaustion, we first treated CAR-T cells with Met or Rap alone, Met+Rap, or without any of these for one week during the expansion, maintained them in a drug-free condition for the next six days, and profiled them by flow cytometry (Supplementary Figure 2A). CAR-T cells without pretreatment showed more effector memory type cells and higher levels of Tim3, PD-1, IFNγ, and TNFα than wt-(non-CAR)T cells (Supplementary Figure 2B). These observations are likely attributed to nonspecific activation signals triggered by CAR, which was mitigated by Rap and Met+Rap treatment (Supplementary Figure 2B).

Next, to investigate their exhaustion resistance, we tested chronic stimulation of CAR-T cells pretreated with the metabolic regulators. After pretreatment with the drugs, to induce exhaustion, CAR-T cells were chronically re-stimulated with CD3/28 under the hypoxic condition for six days following the expansion (9) (Figure 3A). Subsequently, their cytotoxic activity was evaluated against SB28mEGFRvIII cells under hypoxia. CAR-T cells without pretreatment revealed the least level of cytolytic capability, suggesting their exhaustion status. Conversely, those pretreated with Rap and Met+Rap retained their antitumor effect even after chronic stimulation, indicating their better tolerance against exhaustion (Figure 3A).

**Figure 3.**
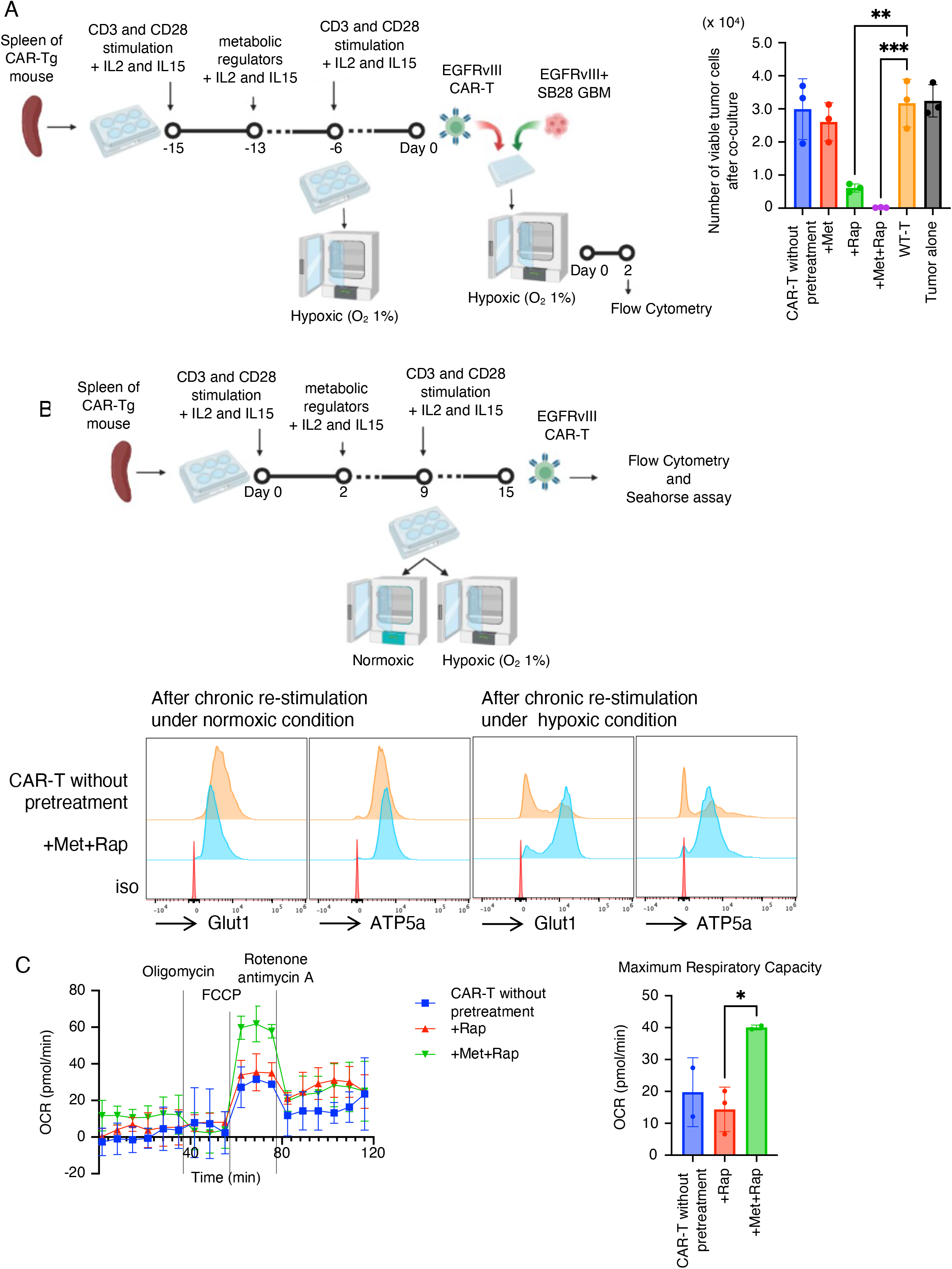
Chronic hypoxic condition-induced exhaustion of CAR-T cells is mitigated by metabolic regulators. (A) Co-culture experimental design following chronic re-stimulation with CD3/28 dynabeads under hypoxic condition (left). Graphs showing the number of live tumor cells after co-culture (right). (B) Experimental design (upper panel). Flow cytometric data for metabolic markers Glut1 and ATP5a in CD8+ CAR-T cells with and without pretreatment with Met+Rap on Day 15 with chronic re-stimulation in normoxic (left) and hypoxic conditions (right). (C) OCR of Day 15 CAR-T cells after the chronic re-stimulation under the hypoxic condition was measured by Seahorse XFe96 analyzer. Data represent the means ± SEM. Maximum Respiratory Capacity was calculated (right). Error bars show the mean with SD. *p < 0.05, **p < 0.01, ***p < 0.001 by one-way ANOVA test followed by Tukey’s multiple comparisons test (A, C).

Previous studies have reported that exhausted T cells exhibit suppressed mitochondrial respiration and glycolysis (9, 18). To investigate whether the metabolic regulators prevent decreases in OXPHOS, we utilized a chronic stimulation model (Figure 3B). After chronic stimulation under normoxia, there was no remarkable difference in the levels of Glut1 and ATP5a between untreated and Met-Rap-pretreated CAR-T cells (Figure 3B). However, chronic stimulation under hypoxia led to reduced GLUT1 and ATP5a in untreated CAR-T cells, suggesting an exhausted status. In contrast, Met+Rapa-pretreated CAR-T cells maintained high levels of both Glut1 and ATP5a even after chronic stimulation under hypoxic condition (Figure 3B). Moreover, Seahorse assay-based Mito stress test demonstrated that the enhanced maximum respiratory capacity by Met+Rap treatment, but not Rap treatment, persisted after the six-day course of chronic hypoxic stimulation (Figure 3C). These data collectively suggest that the potent resistance to exhaustion in CAR-T cells is induced through the improved metabolic status due to the treatment with Met+Rap.

### In vitro treatment of CAR-T cells with Met+Rap improves the survival of mice bearing intracerebral glioma following IV infusion

To determine whether the Met+Rap treatment of CAR-T cells improves their functions in vivo, C57BL/6J mice bearing intracerebral SB28 mEGFRvIII gliomas received a single IV infusion of 1×10^6^ CAR-T cells pretreated with Met or Rap alone, Met+Rap or without any of these (Figure 4A). As shown in Figure 4A, the infusion with Met+Rap-pretreated CAR-T cells resulted in 6/15 mice surviving at 60 days post-tumor inoculation (median survival 49 days), while all the mice receiving non-treated CAR-T cells died by 51 days after tumor implantation (median survival 34 days) (log-rank, p < 0.0001).

**Figure 4.**
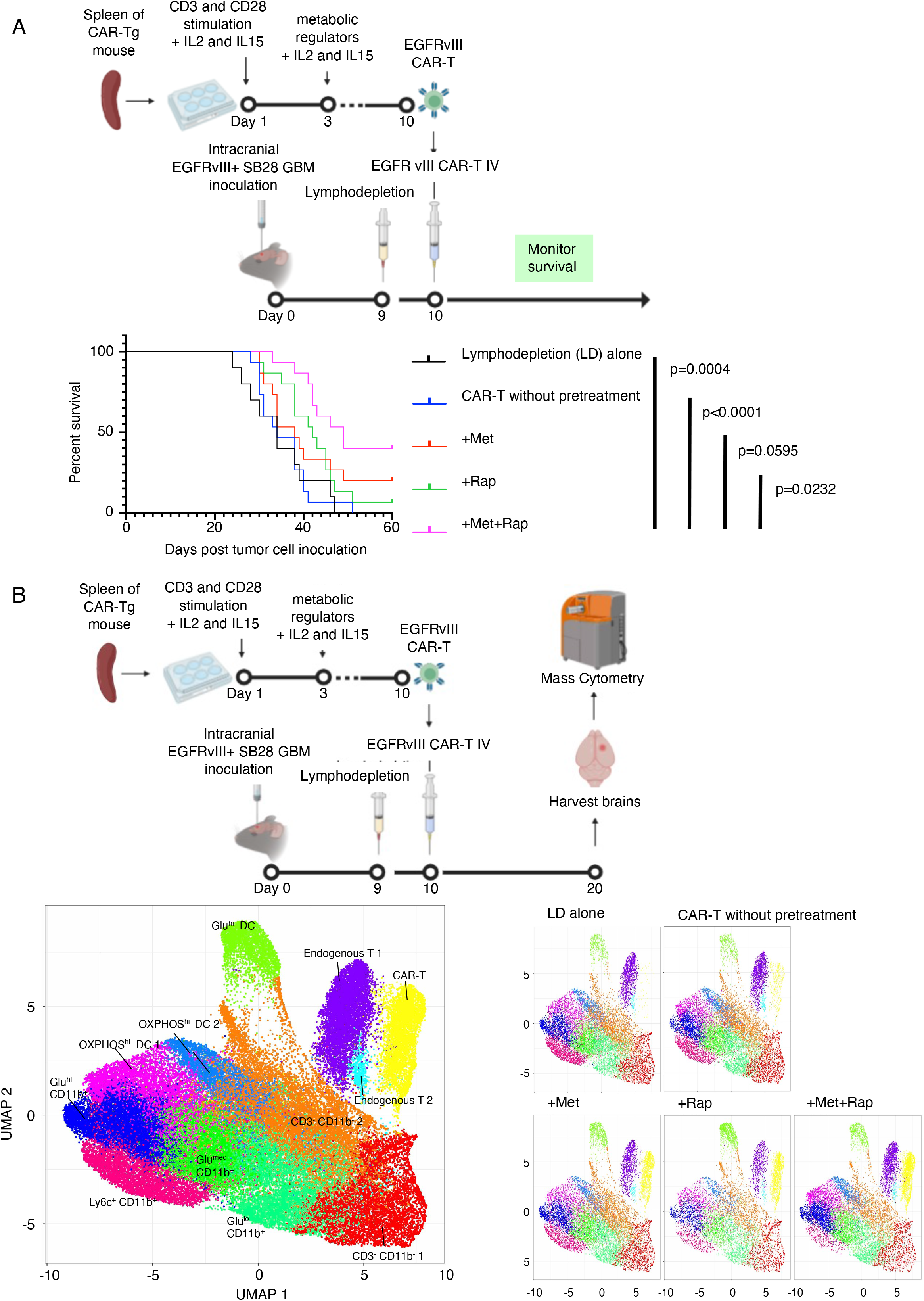
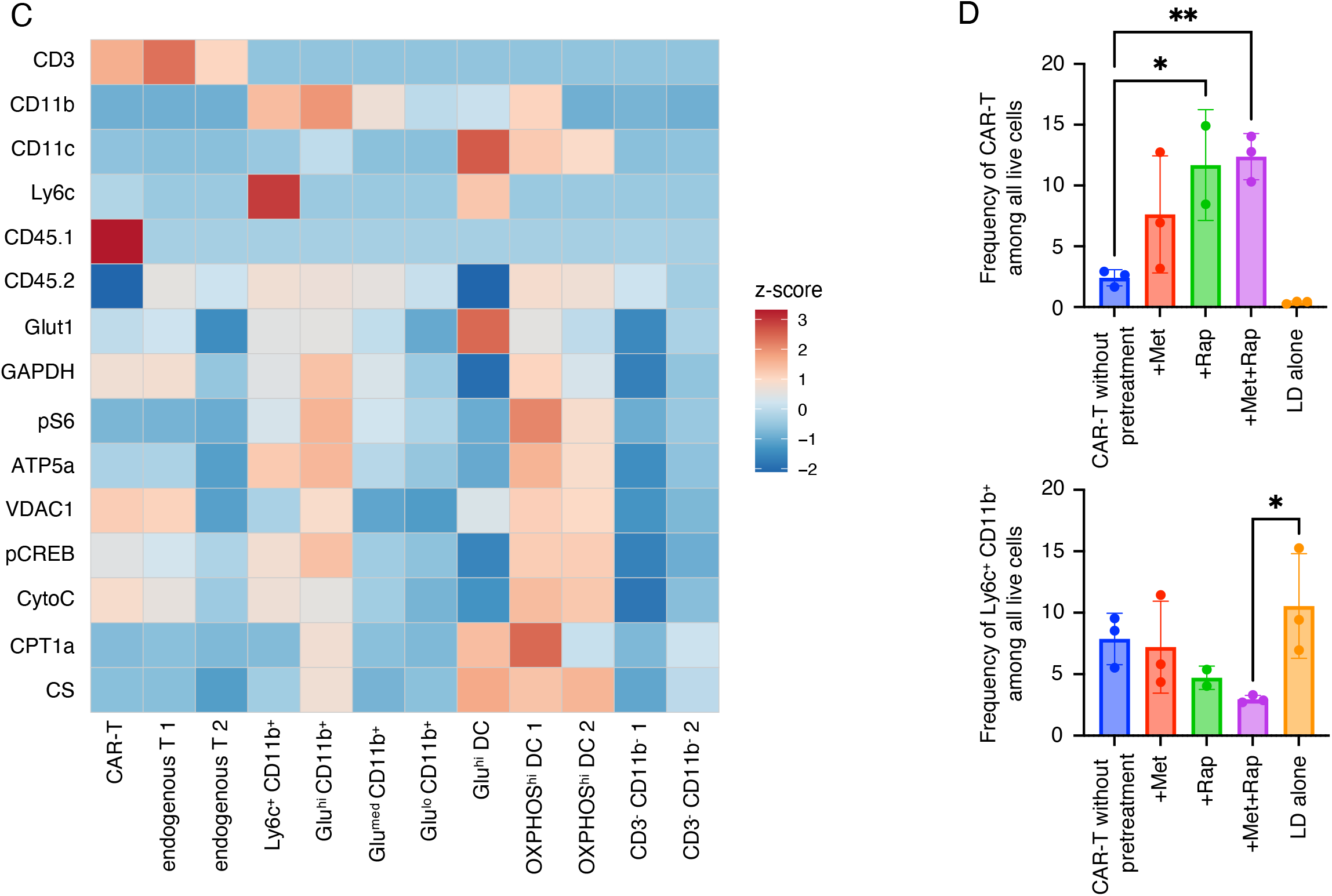
Pretreatment of CAR-T cells with Met+Rap extended the survival of glioma-bearing mice and enhanced their glioma infiltration. (A) Schematic of the treatment protocol for the survival study with the SB28 mEGFRvIII (murine EGFRvIII) glioma (upper panel). Kaplan-Meier curves: LD group (MS = 34 days, n = 10), CAR-T cells without pretreatment (MS = 34 days, n = 15), +Met-pretreated CAR-T (MS = 38 days, n = 15), +Rap-pretreated CAR-T (MS = 42 days, n = 15), and +Met+Rap-pretreated CAR-T (MS = 49 days, n = 15). (B) The design of the mass cytometric analysis (upper panel). Uniform Manifold Approximation and Projection (UMAP) plot of BIL (lower panels). The UMAP in the left penal shows all samples combined, while the panels on the right side show each treatment group separately. (C) Heatmap visualizing the relative expression (z score) of immune cell markers and metabolic markers in each subpopulation. Each cluster was annotated based on the expression status of the markers as indicated in the left panel. Clusters with similar marker expression levels are indicated with labels 1 and 2. (D) Frequencies of SB28 mEGFRvIII-infiltrating CAR-T cells (upper panel) and Ly6c+ CD11b+ monocytic myeloid-derived suppressor cells (MDSCs; lower panel) among the pretreatment types. Error bars show the mean with SD. *P < 0.05; **P < 0.01 by one-way ANOVA analysis followed by Tukey’s multiple comparison test (D).

Previous reports have indicated that pretreatment with IL15 or Rap preserves the stem cell memory phenotype and improves therapeutic outcomes in vivo (19). However, the Met+Rap treatment group demonstrated significantly more extended survival than the Rap alone group (p = 0.0232). In addition to the differences in central memory-dominant T-cell phenotypes, the improved metabolic state of CAR-T cells through Met+Rap pretreatment might have contributed to the observed survival advantages.

To ascertain that Met+Rap treatment is effective across different genders, we further conducted a similar survival study using male mice (Supplementary Figure 3A). Ten mice that were given CAR-T without pretreatment all died by day 41 (Median Survival 37 days), whereas those treated with Met+Rap-pretreated CAR-T were all alive on day 60 (p<0.0001). Additionally, using bioluminescence imaging, we observed a reduction in tumor size in the Met+Rap-pretreated group (Supplementary Figure 3B).

### Mass cytometric analysis reveals the effects of Met+Rap-pretreated CAR-T cells in the glioma microenvironment

To gain an in-depth understanding of how Met+Rap-pretreated CAR-T cells impact the glioma microenvironment, we evaluated the expression of a panel of metabolic markers (20) in the post-infusion glioma tissue using mass cytometry. C57BL/6J mice bearing intracerebral SB28 mEGFR vIII received an IV infusion of 3×10^6^ CD45.1^+^ CAR-T cells pretreated with Met, Rap, Met+Rap, or control non-pretreatment. Ten days after the CAR-T cell administration, we euthanized the mice and profiled brain-infiltrating leukocytes (BILs) by mass cytometry (Figure 4B). We clustered BILs on the uniform manifold approximation and projection (UMAP) plot, grouped into 12 subpopulations, and annotated based on the expression status of lineage markers, glycolysis marker (Glut1, GAPDH, and pS6), and OXPHOS markers (ATP5a, VDAC1, pCREB, CytoC) (Figure 4C). CD45.1^+^ and CD45.2^+^ BILs were glioma-infiltrating CAR-T cells and host C57BL/6J mouse-derived cells, respectively (Figure 4C). We found a significantly higher number of CAR-T cells in the tumor of mice that received Met+Rap-pretreated CAR-T cells than those that received CAR-T cells without pretreatment (Figure 4D, top). Interestingly, the glioma tissues infiltrated by Met+Rap-pretreated CAR-T cells were infiltrated by significantly fewer Ly6C^+^ CD11b^+^ monocytic myeloid-derived suppressor cells (Figure 4D, bottom). These data suggest that Met+Rap-pretreated CAR-T cells have unique abilities to modulate the glioma microenvironment less immunosuppressive.

### Nanostring assay reveals enhanced cytotoxicity and reduced exhaustion of BILs following infusion of Met+Rap-pretreated CAR-T cells

To evaluate the gene expression profile of T cells in the post-treatment tumor microenvironment, we conducted the Nanostring assay following an experimental design similar to mass cytometric analyses. Ten days after the IV CAR-T cell infusion, we extracted the brains, performed FACS sorting to isolate CD3-positive cells, extracted total RNA, and subjected the samples to the Mm Exhaustion panel in the Nanostring assay with biological triplicates (n = 3 mice per group) (Figure 5A). Biological triplicates were conducted for each group, yielding reliable gene expression reproducibility (Figure 5B). The results comparing the group with no pretreatment and the group with Met+Rap pretreatment are depicted in Figure 5C. Remarkably, T cells from the Met+Rap-pretreated group displayed higher expression levels of genes associated with cytotoxicity, hypoxia response, and TCR signaling. In contrast, genes related to mTOR signaling, MAPK signaling, and T cell exhaustion were expressed at higher levels by T cells from the group with no pretreatment. Notably, exhaustion markers, such as *Lag3*, *Pdcd1*, and *Ptger4*, showed a trend toward lower expression levels in the Met+Rap-pretreated group. In contrast, markers of cellular cytotoxicity, such as *Ctsw*, *Prf1*, and *Nkg7*, displayed a trend toward higher expression levels in the Met+Rap-pretreated group (Figure 5D). Although the presence of endogenous T cells limited the detection of significant signal changes, these findings suggest that Met+Rap-pretreated CAR-T cells promote the creation of a more favorable tumor microenvironment.

**Figure 5.**
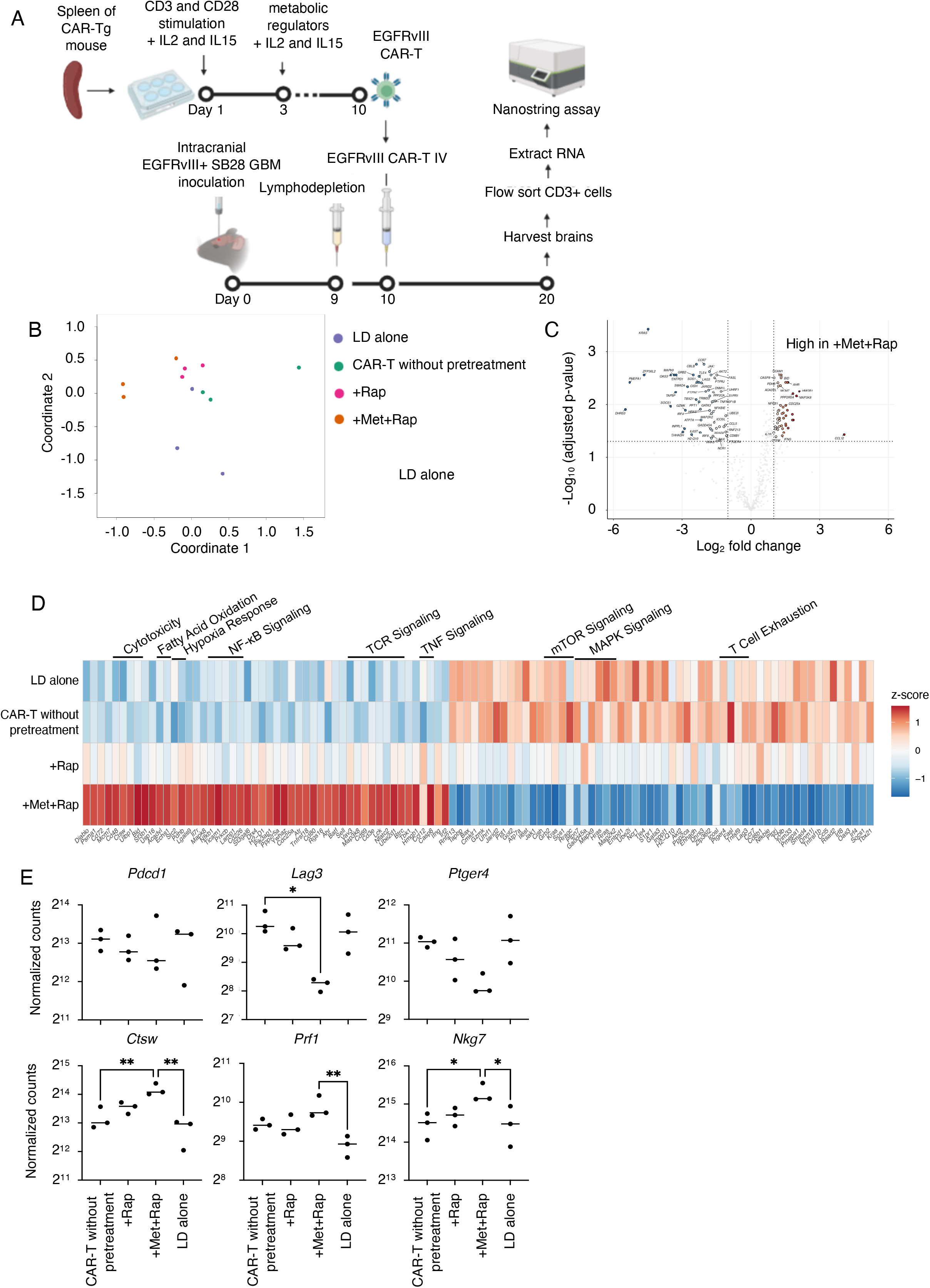
Tumor-infiltrating T cells exhibit enhanced cytotoxic phenotype and lower Lag3 expression in mice that received Met+Rap-pretreated CAR-T cells. (A) The design of the experiment. (B) Multidimensional scaling (MDS) plot colored by the pretreatment types. (C) Comparison of 773 genes between CD8+ CAR-T cells without pretreatment and Met+Rap-pretreated CD8+ CAR-T was summarized in volcano plots. (D) Genes with log2 |fold change|>1.0 and –log10 (adjusted p-value) >1.3 were considered significant. Heatmap visualizing the relative expression (z score) of genes in each subpopulation. (E) Representative gene expression levels obtained from Nanostring analysis, highlighting the most distinctive ones. *p < 0.05, **p < 0.01 by one-way ANOVA test followed by Tukey’s multiple comparisons test (D).

### Met+Rap pretreatment is also effective for human CAR-T cells

A series of murine data collectively showed that the Met+Rap pretreatment could maintain the metabolic status and enhance the therapeutic efficacies of murine CAR-T cells. To determine whether metabolic regulators could enhance the therapeutic effectiveness and metabolic state of human CAR-T cells, we treated human anti-EGFRvIII CAR-T cells with Met and Rap during expansion and co-cultured them with human glioblastoma U87 EGFRvIII cells under hypoxic condition (Figure 6A). The human CAR-T cells were generated by sorting CD8-positive cells from PBMCs derived from healthy donors and then transducing them with lentivirus encoding anti-EGFRvIII CAR (Supplementary Figure 4A). Pretreatment with Rapa or Met+Rap resulted in a slight increase in the central memory population in human CAR-T cells, albeit less pronounced than our observations in mice (Figure 6B). Additionally, as illustrated in Figure 6C, the pretreatment of human CAR-T cells with Met+Rap, but none of the other pretreatments, led to persistent and robust antitumor efficacy even following the challenge with tumor cells five times under the hypoxic condition. These results were consistent across three donors with varying ages, genders, and races (Supplementary Figure 4B). Moreover, the Mito stress test revealed increased SRC in human CAR-T cells pretreated with Met+Rap, as depicted in Figure 6D, suggesting that this therapy could be effective in human CAR-T cells.

**Figure 6.**
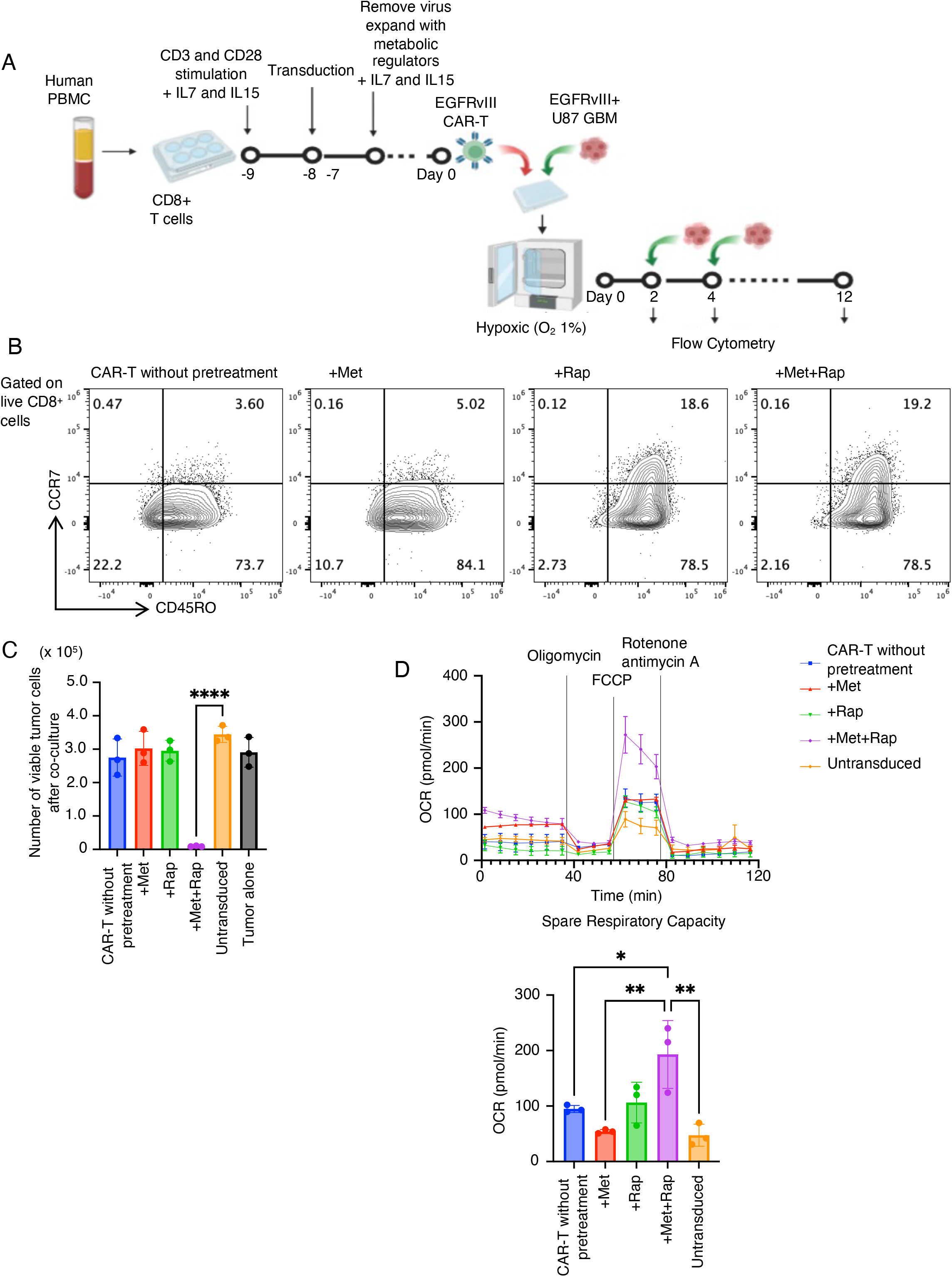
Met+Rap pretreatment enhances the sustained function of human CAR-T cells. (A) Experimental design. (B) Flow cytometric plots for CD45RO and CCR7 in CD8+ CAR-T cells on Day 0. (C) Graphs showing the number of surviving tumor cells after co-culture on Day 10 under hypoxic condition. (D) OCR of CAR-T cells was measured by Seahorse XFe96 analyzer on Day 0 (top). Data represent the means ± SEM. Spare Respiratory Capacity was calculated (bottom). Error bars show the mean with SD. *p < 0.05, ****p < 0.0001 by one-way ANOVA test followed by Tukey’s multiple comparisons test (C-D).

## Discussion

In this study, we grappled with the pivotal challenge of enhancing the efficacy of immunotherapy against GBM, an unrelenting brain tumor known for its resistance to conventional treatments. The intricate microenvironment of GBM, characterized by immune suppression and hypoxic condition, presents significant obstacles to successful immunotherapy outcomes (5). To surmount these challenges, we delved into the potential of metabolic conditioning to enhance the functional capabilities of CAR-T cells within this complex milieu.

In the context of glioma, tumor-induced vascular endothelial damage is recognized for causing microthromboembolization, leading to hypoxic conditions (21). Additionally, our Hypoxiprobe experiments illustrated that immune cells infiltrating intracranial GBM experience a profoundly hypoxic condition compared to those present in subcutaneously transplanted GBM. Therefore, the reduced OXPHOS activity observed in the CAR-T cells within the glioma microenvironment is likely due to tissue hypoxia and the constrained oxygen availability. These observations led us to hypothesize that preconditioning CAR-T cells with metabolic regulators before infusion would enhance their mitochondrial function, making them more resistant to the induction of exhaustion by the GBM’s hypoxic microenvironment.

Our findings lend strong support to the efficacy and feasibility of this approach. Preconditioning CAR-T cells with the Met+Rap combination significantly enhanced their cytotoxic potency and endurance within the hypoxic environment. While one previous study reported direct anti-neoplastic effects of Met+Rap combination treatment in vitro, Met doses used in those studies (10-20 mM) are much higher than those in our research (10 μM) (22). An intriguing study in aging research revealed that mice systemically treated with the Met and Rap combination exhibited extended lifespans (23). In a murine Crohn’s disease model (24), the Met+Rap combination therapy robustly inhibited mTOR signaling and subsequently reduced inflammation. Our investigation demonstrated that low-dose ex vivo treatment with Met effectively activated AMPK, while Rap inhibited mTORC1 and mTORC2, leading to efficient activation of PGC-1ɑ in Met+Rap-pretreated CAR-T cells. Consequently, this resulted in expanded SRC within Met+Rap-pretreated CAR-T cells. These findings were consistent in both murine and human CAR-T cells, highlighting the potential translatability of this approach to human trials.

While prior attempts to enhance T cell metabolism have included both in vitro (25, 26) and in vivo interventions (27), our approach stands out by minimizing the potential of in vivo side effects. This uniqueness rests in our methods solely involving the in vitro expansion phase of CAR-T cell preparation without necessitating in vivo administration of Met or Rap. However, questions linger about the persistency of the improved metabolic states induced in vitro (4). Our study demonstrated that Met+Rap-pretreated CAR-T cells, after chronic stimulation under hypoxic conditions for at least one week, maintained their tumor-killing effects and preserved mitochondrial function, supported by Seahorse assay data.

Most significantly, a single IV infusion of Met+Rap-pretreated CAR-T cells significantly prolonged the survival of mice bearing intracerebral glioma (median overall survival 49 days in Met+Rap group vs. 34 days non-pretreated CAR-T cell group). Furthermore, our in vivo experiments underscored the clinical relevance of Met+Rap-pretreated CAR-T cells. Besides hypoxia, a notable feature of the brain tumor microenvironment is the abundance of MDSCs (28). In our study, we confirmed by mass cytometry that there was a significant reduction in MDSCs in the brains of brain tumor mice treated with Met+Rap-pretreated CAR-T cells. We speculate inflammatory cytokines, including IFNγ, secreted by Met+Rap-pretreated CAR-T cells contributed to the observations (29). Indeed, our nanostring analysis confirmed elevated *Ifng* expression levels when CAR-T cells were pretreated with Met+Rap.

While our study primarily centers on GBM, the implications of our findings may reach far beyond this specific cancer, extending to various solid tumors marked by an immunosuppressive microenvironment. Notably, CAR-T cells have demonstrated limited efficacy in solid tumors compared to their remarkable success in B cell leukemia or lymphoma (30). This diminished efficacy is particularly pronounced in solid tumors characterized by hypoxia (31). By shedding light on this intricate relationship between tumor hypoxic condition and immune response, we open new paths for innovative therapeutic interventions, potentially redefining how we approach GBM and other challenging solid tumors.

In summary, our study advances immunotherapy by introducing metabolic conditioning to amplify CAR-T cell functionality against the intricate challenges posed by GBM. By unraveling our findings’ mechanisms and broader implications, we contribute to ongoing efforts in crafting innovative and effective immunotherapeutic strategies to combat challenging solid tumors.

## Methods

### Mice and Cells

C57BL/6J mice were purchased from Jackson Laboratory (JAX 000664). Mice were approximately 9–10 weeks old during the experiment and maintained under specific pathogen-free conditions at the Animal Facility at UCSF, per an Institutional Animal Care and Use Committee-approved protocol. The C57BL/6-background CD45.1^+^ CAR Transgenic mouse strain (10) and the murine SB28 glioma cell line were established in our lab (32). To create SB28 cells expressing human EGFRvIII, the cells were retrovirally transduced with human EGFRvIII (SB28 hEGFRvIII) as previously described (10). Separately, to generate SB28 cells expressing murine EGFRvIII, we removed the existing green fluorescent protein (GFP) reporter gene from SB28 using CRISPR/Cas9 to prepare the cells for subsequent modifications. Those modified SB28 were transduced with a lentiviral vector encoding mouse EGFRvIII and GFP (kindly provided by Dr. Yi Fan (33)). Post-transduction, GFP-positive cells, which concurrently expressed mEGFRvIII, were isolated by a cell sorter (SB28 mEGFRvIII). Cell lines were cultured in RPMI medium (Gibco) with 10% (v/v) heat-inactivated fetal bovine serum and 1% (v/v) penicillin-streptomycin mixed solution (Gibco, 15070063). Cell lines were free of mycoplasma contamination.

### In vitro T cell cultures

The spleen was harvested to isolate CD3^+^ CAR-T cells from 8-to 12-week-old CAR Transgenic mice. The spleen was minced and treated with ACK (Ammonium-Chloride-Potassium) Lysing buffer for 2 min to lyse the erythrocytes. CD3^+^ CAR-T cells were then purified from lymphocytes using MojoSort™ Mouse CD3 T Cell Isolation Kit according to the manufacturer’s instructions (BioLegend, 480031). CAR-T cells were then activated for 2 days at 1 × 10^6^ cells per 24-well flat-bottomed plates with an equivalent number of CD3/CD28 washed Dynabeads (Gibco, 11453D), 30U/ml hIL-2 (NIH) and 50ng/ml mIL-15 (Peprotech, 21015) in 1 ml of complete RPMI [cRPMI: RPMI 1640 media with 10% FBS, 1% Penicillin-Streptomycin (Gibco, 15070063), 1% HEPES (Gibco, 15630080), 1% Glutamax (Gibco, 35050061), 1% non-essential amino acids (Gibco, 11140076), 1% sodium pyruvate (Gibco, 11360070), 0.5mM 2-Mercaptoethanol (Gibco, 21985023)]. After stimulation, CAR-T was cultured for 7-10 days in a medium containing IL2 and IL15 (30U/ml hIL-2, 50ng/ml mIL-15) as an expansion step. Cell density was monitored daily and maintained in fresh media and cytokines at 0.5-1×10^6^ cells/ml. For metabolic regulator pretreatment, 1×10^6^ cells/ml T cells were treated with metabolic regulators at the doses indicated in Table 1 during the expansion step. The medium was changed to a fresh medium containing fresh metabolic regulators every two days throughout the pretreatment. The compound names and vendor names of the metabolic regulators are listed in Table 1.

### In vitro co-culture

T cells (0.5-1 × 10^5^) were co-cultured with SB28 mEGFRvIII or SB28 hEGFRvIII tumor cells at an effector: target ratio of 1:1 in 96-well flat-bottom plates in triplicate for two days in cRPMI containing IL2 and IL15 (30U/ml hIL-2, 50ng/ml mIL-15). After 2days, cells were collected, washed twice with PBS, and used in FACS analyses. In the FACS data, GFP-positive and Zombie-negative cells were determined to be the surviving tumor cells after co-culture (Supplementary Figure 5). Co-culture under hypoxia was performed at 1% oxygen using a hypoxic incubator (Binder, CD(E6.1)), as it has been reported that the oxygen concentration in a normal brain is 1-5% (34, 35). Furthermore, since gliomas are known to be more hypoxic than normal brains, and several reports have observed gliomas with 1% oxygen concentration in vitro, we also used 1% oxygen in this study (36, 37).

### Mouse therapy model

An aliquot of 5 × 10^3^ SB28 hEGFRvIII or mEGFRvIII cells/mouse was stereotactically injected into the right hemisphere of anesthetized C57BL/6J mice (day 0). Tumor progression was evaluated by luminescence emission on Xenogen IVIS Spectrum after intraperitoneal injection of 1.5 mg of d-luciferin (GoldBio). Before treatment, mice were randomized, so the initial tumor burden in each group was equivalent. A combination of cyclophosphamide (3 – 4 mg/mouse) and fludarabine (1 mg/mouse) was intraperitoneally injected as lymphodepletion (LD) around days 10 – 14. Mice received intravenous (IV) administration with 1×10^6^ CAR-T cells for survival study the day after LD.

### Isolation of brain tumor-infiltrating immune cells

For brain-infiltrating lymphocytes (BILs) analysis, the brain tumor was harvested and sliced into 3-mm pieces with scalpels, followed by digestion with collagenase type IV (Gibco, 17104019) and Deoxyribonuclease I (Worthington Biochemical, LS002007) using a gentleMACS Dissociator (Miltenyi Biotec). Then BILs were isolated using Percoll (Sigma-Aldrich, P1644) as previously described (38).

### Preparation of lentiviral vectors

The pELNS-3C10-CAR was established in our lab (39). HEK293TN cells (5 × 105) were plated on a 6-well plate. At 24 h, pELNS, pAX, and pMD2.G were co-transfected by Fugene HD Transfection Reagent (Promega Corporation, Madison, WI). The supernatant was collected at 48 h, and stored at −80°C.

### Human T-cell isolation

Human healthy donor-derived whole blood was obtained from StemExpress (Cat# LE005F). Then, peripheral blood mononuclear cells were isolated by Ficoll density gradient centrifugation. Human CD8^+^ T cells were isolated by negative selection using the EasySep Human CD8 negative isolation kit (StemCell Technologies, Vancouver, BC).

### Transduction of T-cells by lentiviral vectors

The isolated T-cells (1×10^6^) were resuspended in 1 ml medium per well of a 24-well plate and stimulated with Dynabeads Human T-Activator CD3/CD28 (Thermo Fisher Scientific), IL-7 (Peprotech, 5 ng/ml), and IL-15 (Peprotech, 5 ng/ml) for 24 h. Next, the cells were resuspended in 0.5mL medium mixed with 0.5mL lentiviral pELNS-3C10-CAR vector supernatant and plated on a 24-well plate pre-coated with retronectin and spun at 1000g for 1 h at 32°C. T-cells were then cultured with the lentiviral vector, and the virus was removed after 24 h. Cell lines were cultured in x-vivo medium (Lonza) with 5% (v/v) gamma-irradiated human AB serum (GeminiBio), 10mM N-Acetyl-L-cysteine (Sigma), and 50uM 2-Mercaptoethanol (Gibco, 21985023).

### Flow cytometric analysis

The following monoclonal antibodies (mAbs) were used to detect the respective antigens in the mouse sample: CD45 (30-F11), CD45.2 (104), CD45.1 (A20), CD8 (53-6.7), CD4 (RM4-5), CD3 (17A2), CD11b (M1/70), CD44 (IM7), and CD62L (MEL-14) from BioLegend; Glut1 (EPR3915) from Abcam. ATP5a expression was detected by anti-ATP5a (Abcum, ab110273), followed by secondary staining with goat Anti-Mouse IgG2b heavy chain (ab130790). The following mAbs were used to detect the respective antigens in the human sample: CD45 (HI30), CD8 (HIT3a), CD4 (RPA T4), CD3 (HIT3a), and CD45RO (UCHL1) from BioLegend; CCR7 (3D12) from Invitrogen. Live/dead cell discrimination was performed using Zombie Aqua™ Fixable Viability Kit (BioLegend, 423102). Intracellular staining was performed using a FOXP3 Fixation Kit (eBioscience). All flow cytometry experiments were performed on the Invitrogen Attune NxT (Thermo Fisher Scientific) flow cytometer and analyzed using FlowJo software (FLOWJO, LLC, Ashland, OR, USA). Data were gated on live (Zombie negative) and single cells.

### In vivo hypoxic condition analysis

C57BL/6J mice received a stereotactic injection of 1×10^4^ SB28 mEGFRvIII cells in the right cerebral hemisphere or subcutaneous injection of 1×10^5^ SB28 mEGFRvIII cells mixed 1:1 with Matrigel (Corning, 354230) in the right flank (Day-16). LD was performed on day -1, and 3×10^6^ CAR-T cells were infused IV on day 0. Mice received IV pimonidazole (80 mg/kg, hypoxyprobe, hp11) in PBS 1.5 h before euthanizing on Day 5. Pimonidazole was visualized with anti-pimonidazole antibodies (hypoxyprobe, hp11) using a FOXP3 Fixation Kit.

### Seahorse Metabolic Assays

The oxygen consumption rate (OCR) of treated cells was measured using an Xfe96 Extracellular Flux analyzer (Agilent, Santa Clara, CA, USA). One day before the experiment, the Xfe96 plate was coated with CellTak solution (Cornin, 354240) per the manufacturer’s recommendation. On the day of the experiment, 2-3 x 10^5^ CD8^+^ T cells per well were seeded in the Cell-Tak-coated Xfe96 plate, and the OCR was measured. In the Mito Stress Test, oligomycin (2 uM), carbonyl cyanide p-trifluoromethoxyphenylhydrazone (FCCP) (2 uM), and rotenone/antimycin A (0.5 uM) were injected to obtain maximal and control OCR values. Different parameters from the OCR graph were calculated. Basal oxygen consumption was defined as (the last rate measurement before oligomycin) – (non-mitochondrial respiration). Maximal respiration was defined as (maximum rate measurement after FCCP) – (non-mitochondrial respiration). Spare respiratory capacity (SRC) was calculated by subtracting basal respiration from maximal respiration.

### Western Blotting

CD8^+^ T cells were isolated from in vitro treated CAR-T cells using Mojosort™ Mouse CD8 T Cell Isolation Kit (Biolegend, 480035). After washing cells with PBS twice, CAR-T cells were lysed in M-PER buffer (Thermo Scientific; 78501) with protease inhibitors (MilliporeSigma, 11836170001) and phosphatase inhibitors (MilliporeSigma, 4906845001). After measurement of protein concentration by BCA assay (Thermo Scientific; 23227), a total of 10–20 µg protein was prepared and loaded onto 4 to 20% gradient Mini-PROTEAN TGX Gels (Bio-Rad, 4561093) and electrophoretically separated at 135 V. Proteins in the gels were transferred to the PVDF membrane using the Trans-Blot Turbo Kit by the Trans-Blot Turbo transfer system (Bio-Rad, 1704156). Membranes were then incubated in a blocking buffer (Nacalai Tesque, 13779-01) for 20 minutes at room temperature, followed by incubation with primary antibody overnight at 4 °C in the blocking buffer. After washing, secondary antibody incubations were done at room temperature for 60 min in the blocking buffer. Blots were developed with enhanced ECL Western Blotting Substrate (ThermoFisher, 32106). Primary antibodies recognizing the following proteins were obtained from Cell Signaling Technology: AMPK alpha (2532), phospho-AMPK alpha (2535), p70 S6 kinase (P-S6K) (9202), phospho-P-S6K (Thr389, 9205), pan-Akt (4691), phospho-Akt (Ser473, 9271) S6 Ribosomal Protein (2217), phospho-S6 Ribosomal Protein (Ser240/244), 4E-BP1 (9644) and phosphor-4E-BP1 (Thr37/46, 2855). An antibody recognizing PGC-1ɑ (66369-1-Ig) was obtained from Proteintech. During the detection of PGC-1ɑ, Can Get Signal® immunoreaction enhancer solutions (NKB-101 and NYPBR01, TOYOBO) were used instead of a blocking buffer. The densities of immunoreactive bands were quantified using ImageStudioLight software (LI-COR Biosciences).

### NanoString Gene Expression Analysis

Ten days after CAR-T administration, mice bearing intracerebral SB28 mEGFRvIII tumors were euthanized, and brains were harvested and processed. Mice were treated as groups of 6 mice each, and BILs from 2 mice were pooled as one sample so that three biological replicates were obtained for each group. We further isolated the CD3^+^ cells by FACSAria (BD Biosciences) and used the purified immune cells for RNA extraction. Total RNA was extracted from flow-sorted CD3^+^ cells using the RNEasyPlus Micro Kit (Qiagen, 74034), following the manufacturer’s instructions. RNA purity and integrity were determined with Agilent RNA 6000 Pico Kit (Agilent, 5067-1513) on Bioanalyser 2100 (Agilent).

The cDNA products were amplified for 8 cycles using the nCounter Low RNA Input Kit (NanoString, Seattle, WA, USA) based on the manufacturer’s protocol. Briefly, 4 µL of RNA (0.78ng/ul) was used for the cDNA conversion, and the Mm Exhaustion Low-input Primers were used for the cDNA amplification. The cDNA was then incubated at 95 °C for 2 min. A total of 7.5 µL of the amplified cDNA product was used for the analysis and the nCounter Mm Exhaustion Panel was used. Hybridation reaction was performed for 18 hours at 65°C. After hybridization, samples were loaded on a nCounter SPRINT Cartridge and processed on the nCounter SPRINT™ Profiler. All data normalization was performed by the nSolver 4.0 software (NanoString, Seattle, WA, USA) and ROSALIND^®^ (ROSALIND, Inc, San Diego, CA, USA), as recommended by the manufacturer.

### Data Acquisition for Mass cytometry

Ten days after CAR-T administration, mice bearing intracerebral SB28 mEGFRvIII tumors were euthanized, and brains were harvested and processed. Mice were treated as groups of 12 mice, and BILs from 4 mice were pooled together as one sample so that three samples were obtained for each group. However, mice that received rap-treated CARt had fewer BILs and thus had only two samples. Cryopreserved BILs were thawed 1:10 in thawing media (cRPMI + 25 U/mL Benzonase). Cells were incubated in 5 mM of cisplatin (Cell-ID Cisplatin; Fluidigm), allowing for the distinguishing of live cells. Cells were then fixed with 1.6% PFA and barcoded with Cell-ID 20-Plex Pd Barcoding Kit (Fluidigm). After Fc blocking (TruStain FcX; BioLegend), cells were stained with a metal-conjugated surface antibody cocktail (Table 2). Cells were then permeabilized with a FOXP3 Fixation Kit and stained with an intracellular antibody cocktail (Table 2), followed by resuspension in Iridium intercalator (Cell-ID Intercalator; Fluidigm) solution overnight. Cells were then washed and resuspended in a running buffer consisting of a 1:10 dilution of normalization beads (EQ Four Element Calibration Beads; Fluidigm) in deionized water. Samples were then acquired on the Fluidigm Helios Mass Cytometer, and resultant data were exported to FCS files for further processing.

**Table 2.** A mass cytometry panel for this study.

| Channel | Marker | Vendor | Clone | [Optimal] (µg/ml) |
| --- | --- | --- | --- | --- |
| 89Y | Ter119 | BioLegend | TER-119 | 6 |
| 113In | CD45.2 | BioLegend | 104 | 3 |
| 115In | CD45.1 | BioLegend | A20 | 3 |
| 139La | CD4 | BioLegend | RMA4-5 | 6 |
| 140Ce | CD8 | BioLegend | 53-6.7 | 6 |
| 141Pr | CD38 | BioLegend | 90 | 6 |
| 142Nd | HADHA | Abcam | EPR17940 | 6 |
| 143Nd | Ly6C | BioLegend | HK1.4 | 0.375 |
| 144Nd | pS6 | CST | 2F9 | 3 |
| 145Nd | pCREB | CST | 87G3 | 1.5 |
| 146Nd | NRF1 | Abcam | EPR5554(N) | 3 |
| 147Sm | CD98 | BioLegend | RL388 | 3 |
| 148Ns | Cyclin B1 | BD | GNS1 | 6 |
| 149Sm | CytoC | BioLegend | 6H2.B4 | 0.75 |
| 150Nd | T-bet | BioLegend | 4B10 | 6 |
| 151Eu | pErk | CST | D13 | 6 |
| 152Sm | Ki67 | Thermo/eBioscience | SolA15 | 6 |
| 153Eu | CS | Abcam | ERP8067 | 0.75 |
| 154Sm | mTOR | CST | 7C10 | 6 |
| 155Gd | HIF1a | Thermo/Invitrogen | 16H4L13 | 6 |
| 156Gd | CD62L | R&D | 95218 | 1.5 |
| 157Gd | TCRgd | BioLegend | GL3 | 1.5 |
| 158Gd | ACADM | Abcam | 3B7BH7 | 6 |
| 159Tb | CD127 | BioLegend | A7R34 | 6 |
| 160Gd | ATP5a | Abcam | 7H10BD4F9 | 6 |
| 161Dy | VDAC1 | Abcam | 20B12AF2 | 6 |
| 162Dy | p4EBP1 | CST | 236B4 | 6 |
| 163Dy | ikBa | CST | L35A5 | 3 |
| 164Dy | TCF1/TCF7 | CST | C63D9 | 6 |
| 165Ho | CPT1a | Abcam | 8F6AE9 | 6 |
| 166Er | Glut1 | Abcam | EPR3915 | 0.75 |
| 167Er | CD11c | BioLegend | N418 | 2 |
| 168Er | CD11b | BioLegend | M1/70 | 2 |
| 169Tm | GAPDH | Thermo | 6C5 | 1.5 |
| 170Er | CD3 | BioLegend | 17A2 | 3 |
| 171Yb | CD103 | BioLegend | 2E7 | 2 |
| 172Yb | KLRG1 | BioLegend | 2F1 | 0.75 |
| 173Yb | PD1 | BioLegend | 29F.1A12 | 0.75 |
| 174Yb | CD69 | R&D | polyclonal | 6 |
| 175Lu | CD44 | BioLegend | IM7 | 0.375 |
| 176Yb | CD27 | BioLegend | LG.3A10 | 1.5 |
| 209Bi | Granzyme B | BioLegend | QA16A02 | 0.75 |

### Processing of mass cytometry data

Raw FCS files were processed by the normalizer function provided by the Parker Institute of Cancer Immunotherapy Premessa package on R Studio. Normalization bead removal and de-barcoding were performed on the same platform. Each immune subpopulation, such as CD3^+^T cells, was gated and exported using FlowJo software. These exported files were then uploaded to the Cytofkit2 package, where immune cells were subjected to dimension reductional algorithm t-SNE or UMAP for visualization in 2D space and clustered using FlowSOM. Standard settings were utilized (with k = 15) The cells in each cluster were then phenotyped and analyzed using z score–normalized marker expression and population data, respectively. All analytic outputs were generated on R Studio and Prism. Statistics A comparison of survival curves between the two groups was tested by the Log-rank test. The variations of data were evaluated as the means ± standard error of the mean (SEM). R software (Version 4.2.1) and Prism software (Version 9; GraphPad Software) were used for data management and statistical analyses. Significance levels were set at 0.05 for all tests.

### Study approval

All animal studies strictly followed UCSF institutional guidelines (Laboratory Animal Resource Center approval no. ♙N185402-02).

## Data and materials availability

All data associated with this study are available in the main text or the supplementary materials.

## Author contributions

RH and HO designed the research. RH, KK, AY, TC, SP, and PC conducted experiments. RH, KK, TN, and LSL were involved in the analysis and interpretation of data. MHS and HO provided overall study supervision. RH, KK, AY, TN, LSL, MHS, and HO wrote the paper.

## Acknowledgments

We would like to express our gratitude to the individuals and facilities that have contributed to this study: the UCSF Parnassus Flow Cytometry Core provided invaluable mass cytometry services, including access to the CyTOF2 Charmander; the UCSF Core Facilities: Laboratory for Cell Analysis offered essential flow cytometry services and Genome services; the Gladstone Institutes Histology and Light Microscopy Core, along with Dr. Ken Nakamura, graciously permitted the use of the Xfe96 Extracellular Flux Analyzer. The experimental designs in Figures are generated with BioRender (https://biorender.com/). Our research received generous support from NIH grant 1R35NS105068, awarded to H. Okada. We also acknowledge the financial support of the Panattoni Family Foundation, the Parker Institute for Cancer Immunotherapy, both attributed to H. Okada, and the JSPS Overseas Research Fellowships granted to R. Hatae.

## Abbreviations

CAR-T, chimeric antigen receptor-transduced T cells; IV, intravenous; IC, intracranial tumor models; SC, subcutaneous tumor models; MFI, Mean Fluorescence Intensity; Met, Metformin; Rap, Rapamycin; Met+Rap, combination therapy of Metformin and Rapamycin; 2DG, 2-Deoxy-D-glucose; DCA, Dichloroacetic acid; WT, wild-type; LD, lymphodepletion; p-, phospho-; PGC-1α, PPAR-gamma coactivator 1α; SEM, standard error of mean; MS, median survival; Glut1, glucose transporter 1; ATP5a, ATP synthase; VDAC1, voltage-dependent ion channel 1; pCREB, phosphorylation of cAMP response element binding protein; CytoC, cytochrome C; CPT1a, carnitine palmitoyltransferase 1A; CS, citrate synthase; DC, dendritic cells; UT, untransduced cells.

