## Supplementary Figure 1 for "Enhancing CAR-T Cell Metabolism to Overcome Hypoxic Conditions in the Brain Tumor Microenvironment"

### Slide 1
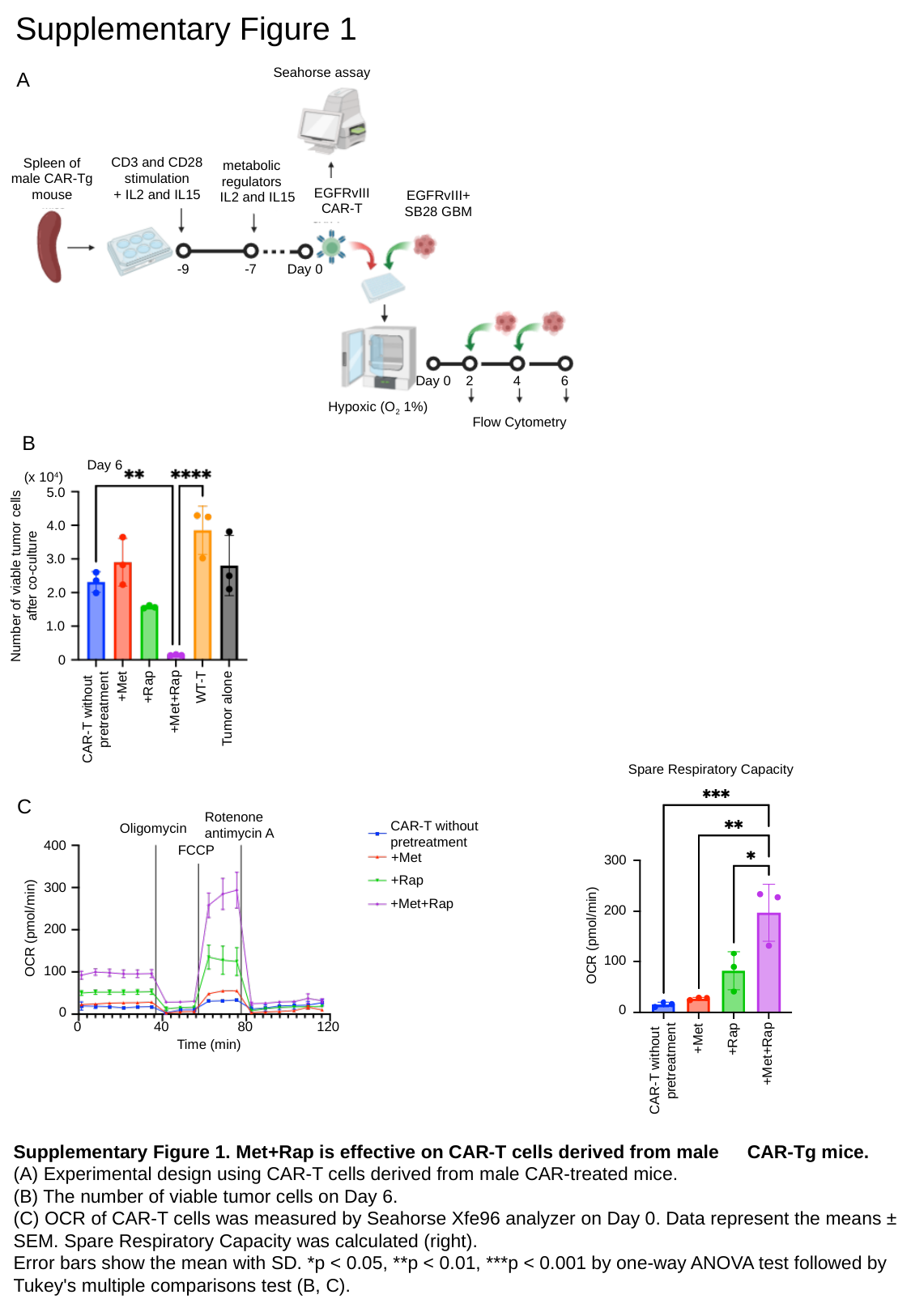

Supplementary Figure 1
Seahorse assay
A
CD3 and CD28 stimulation
+ IL2 and IL15
Spleen of
male CAR-Tg mouse
metabolic
regulators
+ IL2 and IL15
EGFRvIII
CAR-T
EGFRvIII+
SB28 GBM
-9
-7
Day 0
Day 0
2
4
6
Hypoxic (O_2_ 1%)
Hypoxic (O_2_ 1%)
Flow Cytometry
B
Day 6
(x 10^4^)
5.0
4.0
3.0
Number of viable tumor cells
after co-culture
2.0
1.0
0
+Met
WT-T
+Rap
+Met+Rap
Tumor alone
CAR-T without
pretreatment
Spare Respiratory Capacity
C
Rotenone
antimycin A
CAR-T without
pretreatment
Oligomycin
400
FCCP
+Met
300
+Rap
300
+Met+Rap
200
200
OCR (pmol/min)
OCR (pmol/min)
100
100
0
0
120
0
40
80
+Met
+Rap
Time (min)
+Met+Rap
CAR-T without
pretreatment
Supplementary Figure 1. Met+Rap is effective on CAR-T cells derived from male　CAR-Tg mice.
(A) Experimental design using CAR-T cells derived from male CAR-treated mice.
(B) The number of viable tumor cells on Day 6.
(C) OCR of CAR-T cells was measured by Seahorse Xfe96 analyzer on Day 0. Data represent the means ± SEM. Spare Respiratory Capacity was calculated (right).
Error bars show the mean with SD. *p < 0.05, **p < 0.01, ***p < 0.001 by one-way ANOVA test followed by Tukey's multiple comparisons test (B, C).
