## Supplementary Figure 2 for "Enhancing CAR-T Cell Metabolism to Overcome Hypoxic Conditions in the Brain Tumor Microenvironment"

### Slide 1
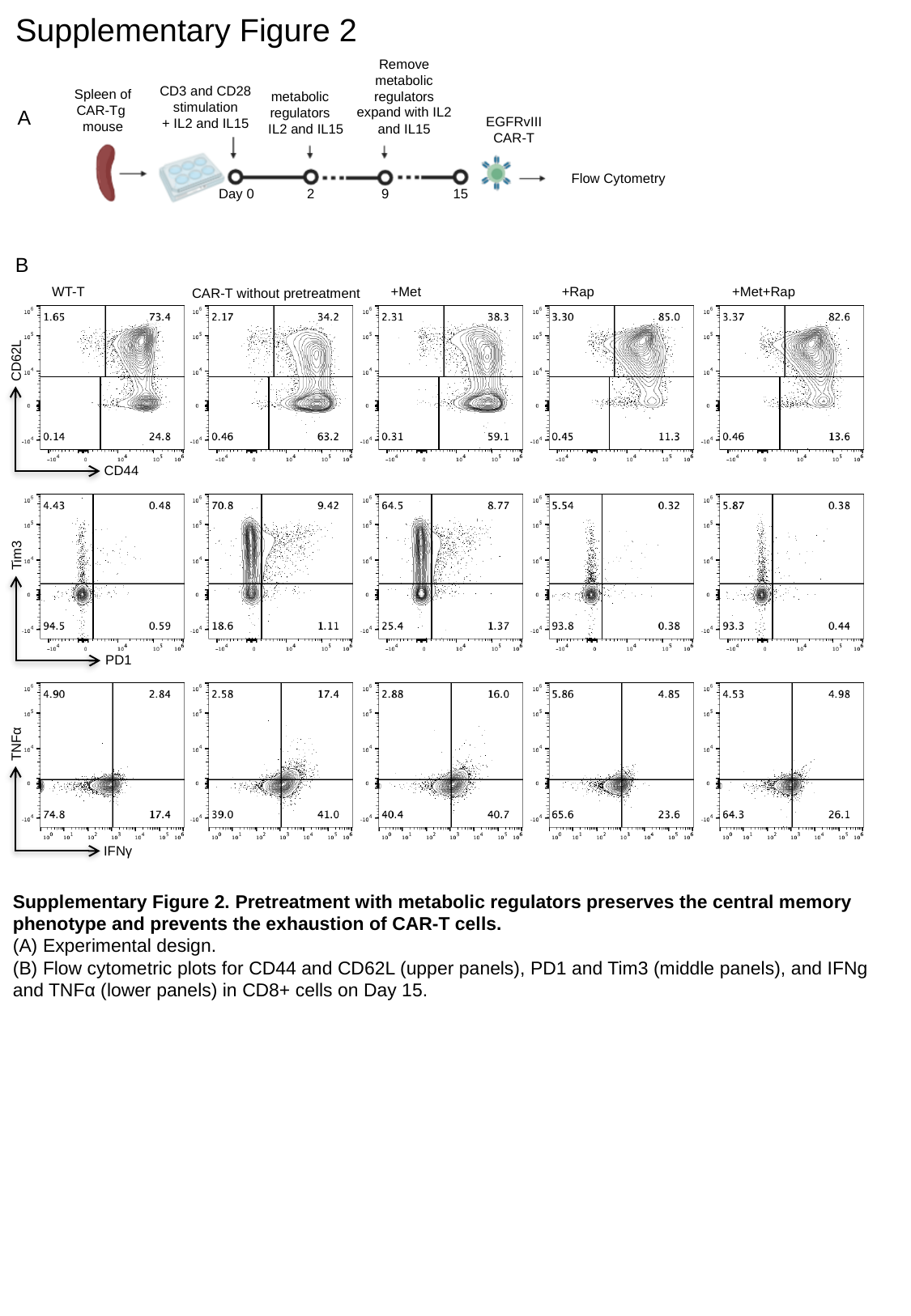

Supplementary Figure 2
Remove metabolic regulators
expand with IL2 and IL15
CD3 and CD28 stimulation
+ IL2 and IL15
Spleen of
CAR-Tg
mouse
metabolic
regulators
+ IL2 and IL15
A
EGFRvIII
CAR-T
Flow Cytometry
2
Day 0
9
15
B
WT-T
+Met
+Rap
+Met+Rap
CAR-T without pretreatment
CD62L
CD44
Tim3
PD1
TNFα
IFNγ
Supplementary Figure 2. Pretreatment with metabolic regulators preserves the central memory phenotype and prevents the exhaustion of CAR-T cells.
(A) Experimental design.
(B) Flow cytometric plots for CD44 and CD62L (upper panels), PD1 and Tim3 (middle panels), and IFNg and TNFα (lower panels) in CD8+ cells on Day 15.
