## Supplementary Figure 3 for "Enhancing CAR-T Cell Metabolism to Overcome Hypoxic Conditions in the Brain Tumor Microenvironment"

### Slide 1
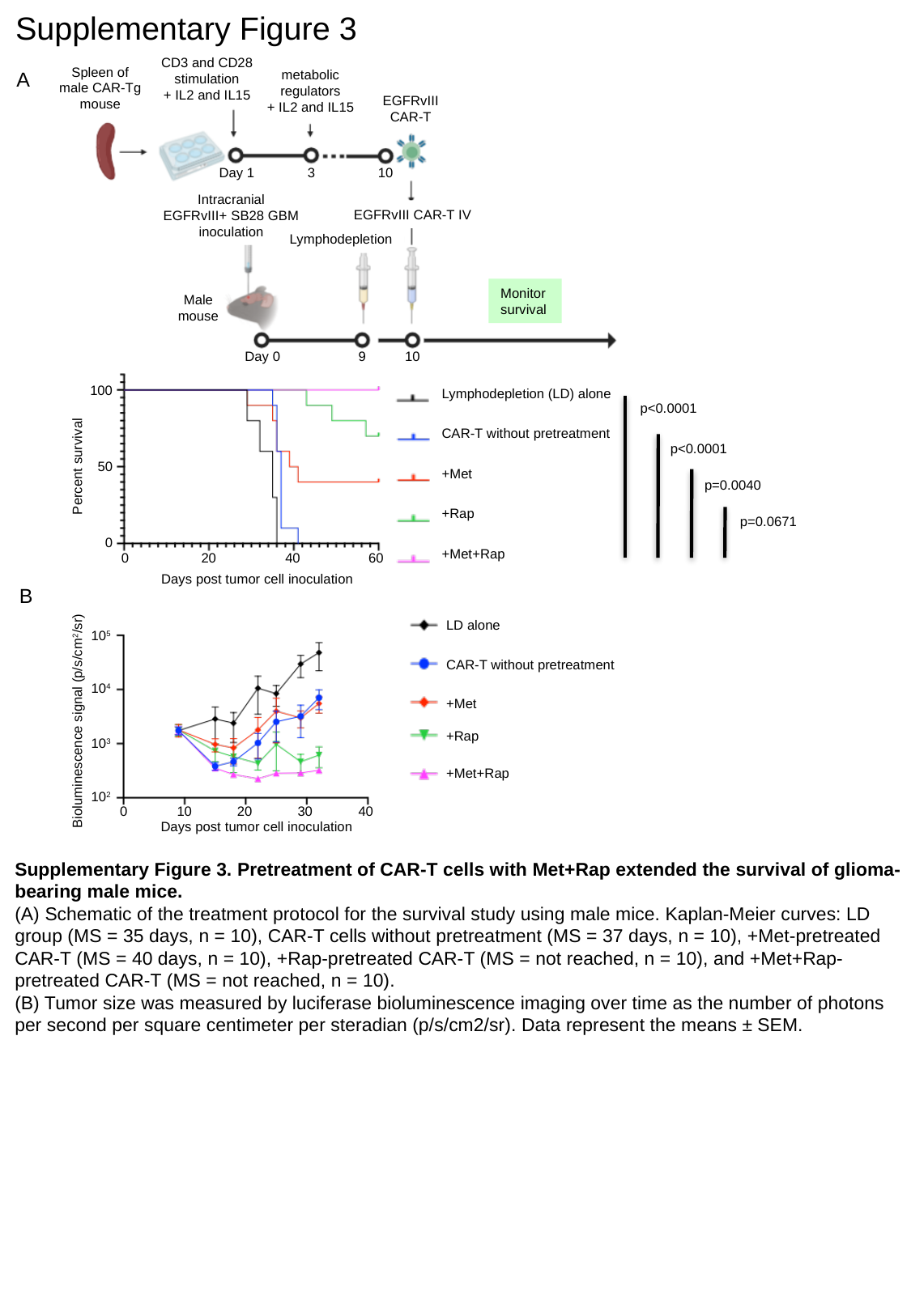

Supplementary Figure 3
CD3 and CD28 stimulation
+ IL2 and IL15
Spleen of
male CAR-Tg mouse
A
metabolic
regulators
+ IL2 and IL15
Monitor
survival
EGFRvIII
CAR-T
Day 1
3
10
Intracranial
EGFRvIII+ SB28 GBM
inoculation
EGFRvIII CAR-T IV
Lymphodepletion
Male
mouse
Lymphodepletion (LD) alone
CAR-T without pretreatment
+Met
+Rap
+Met+Rap
Day 0
9
10
100
p<0.0001
p<0.0001
p=0.0040
p=0.0671
50
Percent survival
0
0
20
40
60
Days post tumor cell inoculation
B
LD alone
CAR-T without pretreatment
+Met
+Rap
+Met+Rap
10^5^
10^4^
Bioluminescence signal (p/s/cm^2^/sr)
10^3^
10^2^
0
20
40
10
30
Days post tumor cell inoculation
Supplementary Figure 3. Pretreatment of CAR-T cells with Met+Rap extended the survival of glioma-bearing male mice.
(A) Schematic of the treatment protocol for the survival study using male mice. Kaplan-Meier curves: LD group (MS = 35 days, n = 10), CAR-T cells without pretreatment (MS = 37 days, n = 10), +Met-pretreated CAR-T (MS = 40 days, n = 10), +Rap-pretreated CAR-T (MS = not reached, n = 10), and +Met+Rap-pretreated CAR-T (MS = not reached, n = 10).
(B) Tumor size was measured by luciferase bioluminescence imaging over time as the number of photons per second per square centimeter per steradian (p/s/cm2/sr). Data represent the means ± SEM.
