## Supplementary Figure 4 for "Enhancing CAR-T Cell Metabolism to Overcome Hypoxic Conditions in the Brain Tumor Microenvironment"

### Slide 1
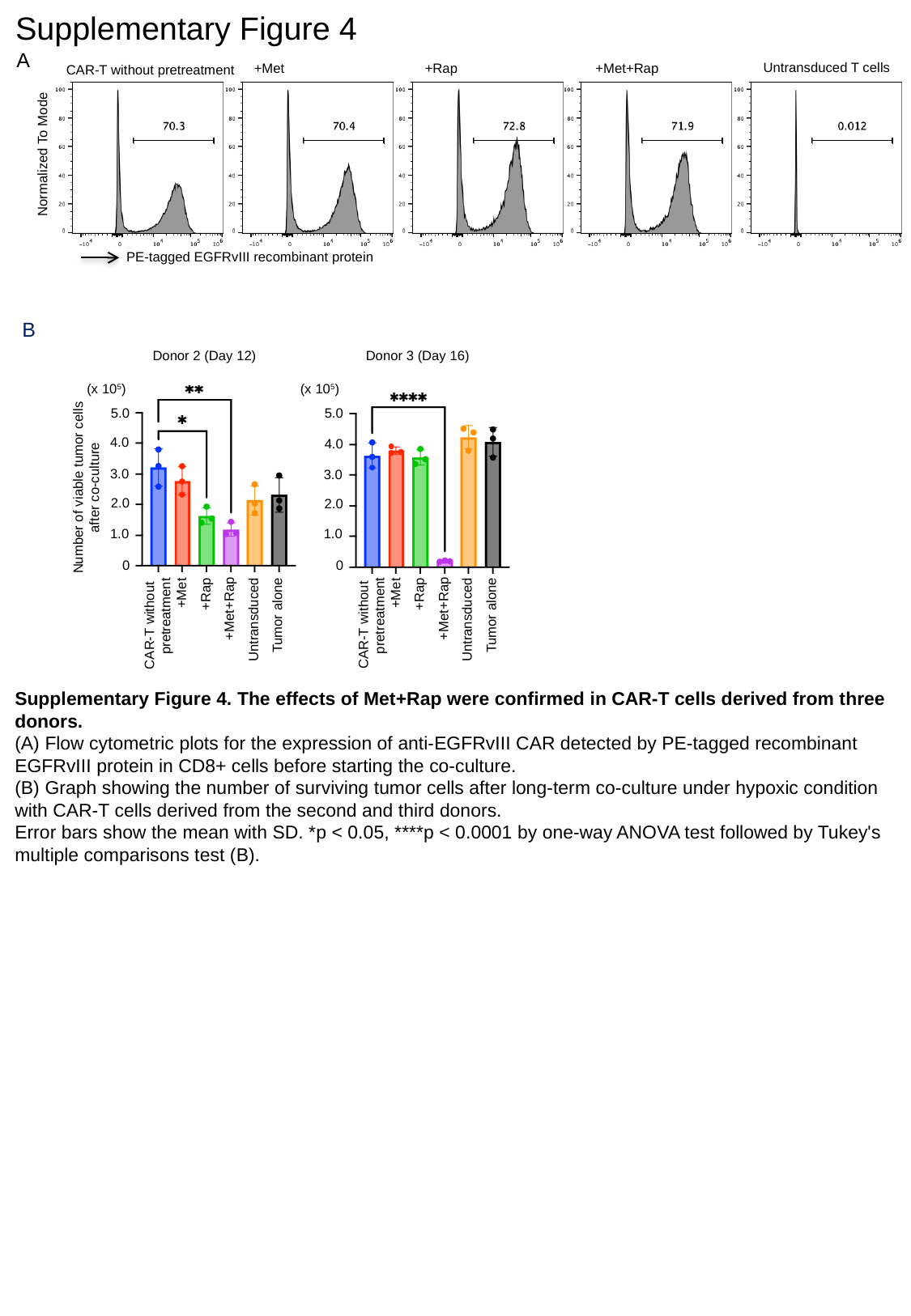

Supplementary Figure 4
A
Untransduced T cells
+Met
+Rap
+Met+Rap
CAR-T without pretreatment
Normalized To Mode
PE-tagged EGFRvIII recombinant protein
B
Donor 2 (Day 12)
Donor 3 (Day 16)
(x 10^5^)
(x 10^5^)
5.0
5.0
4.0
4.0
3.0
3.0
Number of viable tumor cells
after co-culture
2.0
2.0
1.0
1.0
0
0
+Met
+Met
+Rap
+Rap
+Met+Rap
+Met+Rap
CAR-T without
pretreatment
Tumor alone
Tumor alone
CAR-T without
pretreatment
Untransduced
Untransduced
Supplementary Figure 4. The effects of Met+Rap were confirmed in CAR-T cells derived from three donors.
(A) Flow cytometric plots for the expression of anti-EGFRvIII CAR detected by PE-tagged recombinant EGFRvIII protein in CD8+ cells before starting the co-culture.
(B) Graph showing the number of surviving tumor cells after long-term co-culture under hypoxic condition with CAR-T cells derived from the second and third donors.
Error bars show the mean with SD. *p < 0.05, ****p < 0.0001 by one-way ANOVA test followed by Tukey's multiple comparisons test (B).
