## Supplementary Figure 5 for "Enhancing CAR-T Cell Metabolism to Overcome Hypoxic Conditions in the Brain Tumor Microenvironment"

### Slide 1
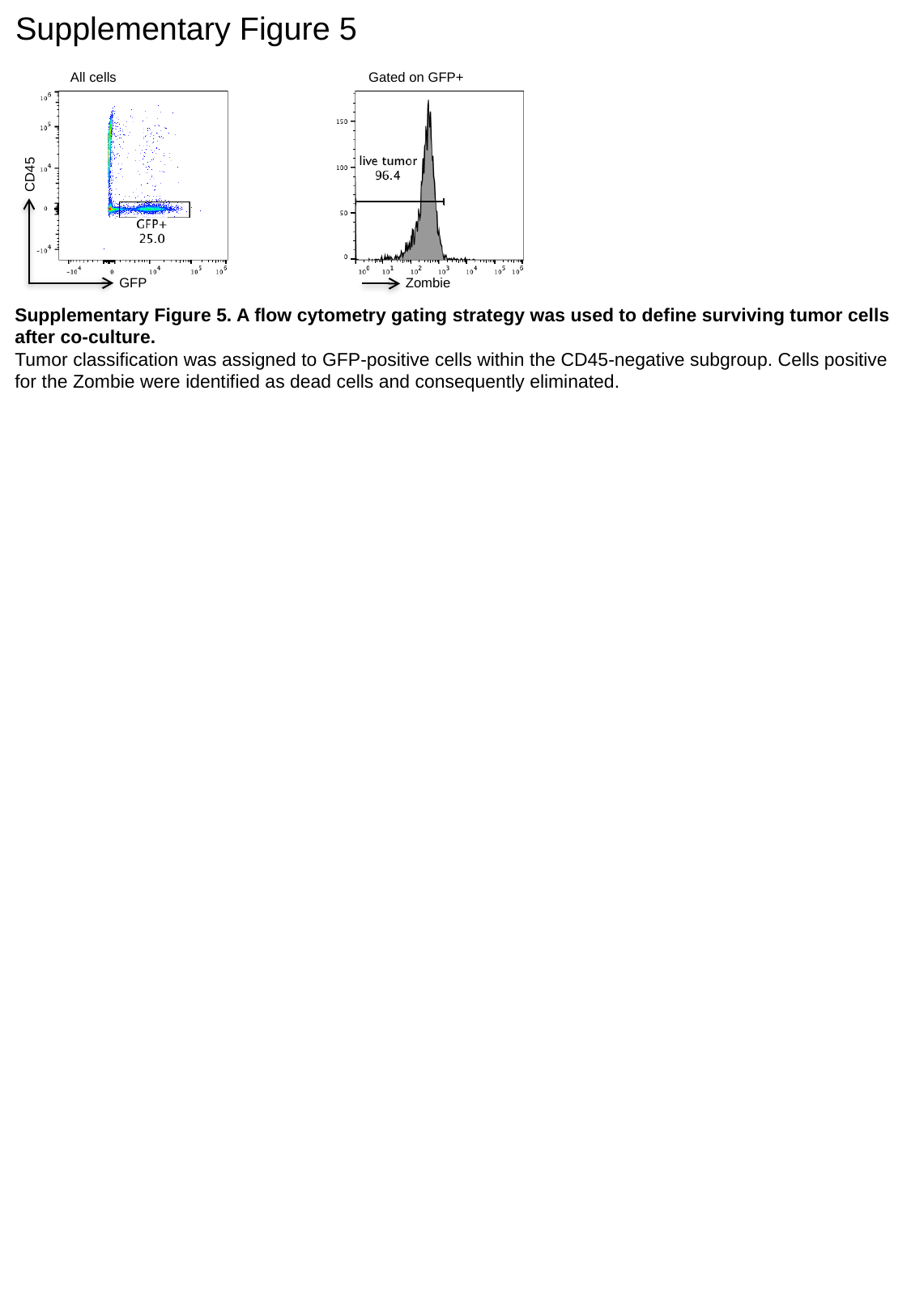

Supplementary Figure 5
All cells
Gated on GFP+
CD45
GFP
Zombie
Supplementary Figure 5. A flow cytometry gating strategy was used to define surviving tumor cells after co-culture.
Tumor classification was assigned to GFP-positive cells within the CD45-negative subgroup. Cells positive for the Zombie were identified as dead cells and consequently eliminated.
